# MicroRNA-17∼92 drives metabolic programming of virus-specific effector CD4 and CD8 T cell responses

**DOI:** 10.64898/2026.02.01.703165

**Authors:** Surojit Sarkar, Yevgeniy Yuzefpolskiy, Florian M. Baumann, Surojit Sarkar, Vandana Kalia

## Abstract

The miR-17∼92 microRNA cluster drives oncogenesis by inducing proliferation and survival of cancer cells. A similar pro-proliferative role of miR-17∼92 has also been identified in pathogen- and tumor-reactive T cells. However, the metabolic underpinnings of miR-17∼92-drivenT cell expansion and effector differentiation remain undefined. Using a murine model of conditional miR-17∼92 deletion or constitutive overexpression in viral antigen-specific CD4 or CD8 T cells, here we show that miR-17∼92 drives terminal differentiation of CD4 and CD8 T cells through sustained activation of mTOR and glycolytic bioenergetics pathways. Constitutively increased expression of miR-17∼92 led to a universal increase in the numbers of antigen-specific T follicular helper (T_FH_), T helper 1 (T_H1_) and cytotoxic T lymphocyte (CTL) effector subsets. During early stages of T cell activation, miR-17∼92 overexpression was associated with increased glucose uptake and heightened glycolytic and oxidative metabolism to sustain increased proliferation in both CD4 and CD8 T cells. However, prolonged overexpression of miR-17∼92 led to a loss of polyfunctionality and compromised metabolic fitness (glycolysis and mitochondrial respiration) by the peak of the effector responses. These studies establish miR-17∼92 as a critical driver of T cell proliferation and effector differentiation through metabolic regulation of both effector CTL and CD4 T cell subsets (T_FH_ and T_H1_) that are critical for combating viral infections. These studies form the basis for future manipulation of miR-17∼92 gene/metabolic regulatory network to commandeer T cell immunity during infection, vaccination or cancer immunotherapy.

## INTRODUCTION

MicroRNAs (miRNAs) are short nucleotide sequences that serve as potent regulators of protein expression at the mRNA level (1). Many evolutionarily conserved miRNAs are important in regulating the development, breadth, and function of T cells (2–9). One such cluster is miR-17∼92, an oncogenic miRNA cluster overexpressed in multiple tumor types (10–13). miR-17∼92 induction in tumor cells is typically associated with increased cMyc expression, which is implicated in sustaining exponential expansion through increased glucose consumption, lactic acid fermentation, and glutamine degradation (14–16).

The miR-17∼92 cluster is upregulated in T cells upon antigen priming, and over-expression of the miR-17∼92 cluster in somatic murine cells results in systemic lymphoproliferation and autoimmunity, driven in part by over-activation of CD4 T cells, even under homeostatic conditions (17). Studies from our lab and others (18, 19), have established that miR-17∼92 is an important mediator of antiviral CD8 T cell expansion and differentiation; over-expression of miR-17∼92 leads to increased expansion and terminal differentiation of antigen-specific effector CD8 T cells (CTL), whereas loss of miR-17∼92 expression in antigen-specific CD8 T cells leads to reduced expansion and preservation of memory-precursor properties. In CD4 T cell responses, miR-17∼92 has been shown to inhibit Treg cell differentiation, while promoting T follicular helper (T_FH_) and T_H1_ effector responses (20–24). T_FH_ and T_H1_ responses are typically induced following intracellular infection and exert critical roles in primary as well as secondary immune responses. Consistent with the well-established role of T_FH_ cells in promoting antibody responses, miR-17∼92-dependent induction of T_FH_ cells was associated with increased germinal centers and antibody responses (6, 23). Likewise, consistent with the role of T_H1_ cells in promoting CTL responses through cytokine mediators and licensing of antigen presenting cells, miR-17∼92 promoted T_H1_ responses in addition to CTL responses (20, 22).

T cell priming and expansion is associated with conversion from a resting metabolic state to an activated state of energy production and nutrient formation utilizing the tumor-associated process of aerobic glycolysis (Warburg effect) (25). Based on the strong association of metabolic state with T cell fates (26–28), the goal of this study was to determine whether miR-17∼92 dictated terminal differentiation of effector T cells through modulation of their metabolic activities. We engaged established murine models of conditional transgene overexpression (CTg) or deletion (conditional knock-out, CKO) of miR-17∼92 in activated antigen-specific T cells to specifically query the role of miR-17∼92 in metabolic regulation of CTL, T_FH_ and T_H1_ differentiation during acute LCMV infection. Using conditional gene regulation models that alter miR-17∼92 expression specifically in post-thymic, mature peripheral T cells immediately after activation, our studies uncover a common program of increased oxidative phosphorylation, aerobic glycolysis, proliferation and terminal differentiation by miR-17∼92 in CD8, T_FH_ and T_H1_ cells alike. This is mediated by sustained activation of the PI3K/Akt/mTOR axis under conditions of increased miR-17∼92 expression. These studies reveal a novel role for miR-17∼92 as a metabolic modulator of CTL and T_H_ cell differentiation following viral infection and emphasize its use as a potential target to manipulate the immune outcomes in future vaccine designs and immunotherapies.

## MATERIALS AND METHODS

### Mice

C57BL/6 mice were purchased from the Jackson Laboratory (Bar Harbor, ME, USA). Thy1.1+ P14 mice bearing the H-2D^b^GP33 epitope-specific TCR were fully backcrossed to C57BL/6 mice and were maintained in our animal colony. CD4 TCR-transgenic Smarta mice, which have CD4 T cells specific for the GP67-77 (KGVYQFKSV) epitope of LCMV, were also maintained in our colony. Floxed mice for conditional deletion (Mir7-92^tm1.1Tyj^/J) or constitutive expression (C57BL/6-Gt(ROSA)26^Sortm3(CAG-MIR17-92,-EGFP)Rsky^/J) of miR-17∼92 were purchased from the Jackson Laboratory (Catalog numbers 008458 and 008517, respectively). These were each crossed with Granzyme B-Cre mice (a generous gift from Dr. J. Jacob) to generate conditional knockout (CKO) or conditional transgenic (CTg) mice in which Cre recombinase is expressed under control of the human granzyme B (GzmB) promoter (29). Antigen-specific T cells rapidly upregulate GzmB following antigen priming causing over-expression or deletion of miR-17∼92. Mice were sacrificed on day 8 post-infection and single cell suspensions of spleen, inguinal lymph nodes, lungs, or PBLs from mice were prepared and direct *ex vivo* staining was carried out as described previously (30–32). All animals were used in accordance with University Institutional Animal Care and Use Committee guidelines.

### Virus and infection

Armstrong strain of LCMV was propagated, titered, and used as previously described (30–33). Mice were injected intraperitoneally with 2×10^5^ pfu LCMV_Arm_.

### *In vitro* stimulation

Bulk naïve CD4 and CD8 T cells were sorted from naïve mice on CD4+ or CD8+, and CD44^Lo^. Cells were stimulated *in vitro* for 24-96 hours on αCD3+αCD28 coated plates at 37 °C and 5% CO_2_ with 10% FBS RPMI. To assess the role of miR17∼92 in mTOR mediated signaling and metabolism, T cells were treated with Rapamycin at 20ng/mL starting day 1 post-activation.

### Seahorse Assay

Oxygen consumption and extracellular acidification were measured using the XFe96 well Seahorse Analyzer (Agilent) as previously described (31). Cells were sorted on Thy1.2, CD4, CD8 and CD44 expression from mice at indicated days post-infection and adhered to plates using poly-L-lysine at 100K (*in vitro* activated) or 120K (direct *ex vivo* acquired) cells per well. Mitostress test was performed following 24hr or 4 days post-activation, and 8 days post-infection in the presence of 10mM glucose (Agilent) with an addition of 2-Deoxyglucose (20mM) to obtain the zero point value of extracellular acidification.

### Flow cytometry

All antibodies were purchased from Biolegend (San Diego, CA, USA) unless stated otherwise: CD4 (RM4-5), CD44 (IM7), IFN-γ (XMG1.2), BrdU (PRB-1 Invitrogen), Ly6c (AL-21 BD Biosciences), CXCR5 (L138D7), T-bet (4B10 Santa Cruz), Bcl-6 (IG191E/A8), PD-1 (RMP1-30), Thy1.1 (OX-7), Thy1.2 (30-H12), CD8 (53-6.7), granzyme B (GB11). MHC class II tetramer was obtained the National Institute of Health Tetramer Core Facility (Atlanta, GA). Cells were stained for surface or intracellular proteins and cytokines as previously described (31). For analysis of intracellular cytokines, 2×10^6^ lymphocytes were stimulated with 0.2μg/ml GP61, GP33, or GP276 peptides in the presence of brefeldin A for 5h. Flow cytometric analysis was performed on LSRII Fortessa (BD Biosciences, San Jose, CA). To assess phosphosignaling, cells were stimulated for 15 minutes in the presence of GP33 peptide and then fixed using PFA. Cells were permeabilized using MeOH and then stained for pS6 (BD Biosciences N7-548).

### Statistical analysis

Unpaired Student’s t-test was used as indicated to evaluate differences between sample means of two groups. More than two means were compared using a one-way ANOVA with a post-hoc Tukey’s post-test analysis. All statistical analyses were performed using Prism 5 and P values of statistical significance are depicted by asterisk per the Michelin guide scale: * (P ≤ 0.05), ** (P ≤ 0.01), *** (P ≤ 0.001) and (P > 0.05) was considered not significant (ns). GSEA analysis was performed on previously deposited microarray data under GEO accession number GSE42760, and GSE34218 using Gene Sets: Mootha Mitochondrial, Module 152 (genes utilized in oxidative phosphorylation) and Mitosis.

## RESULTS

### miR-17∼92 critically regulates metabolic activity of effector CD4 and T cells

Differentiation from naïve-to-effector T cell state is characterized by increased glucose uptake and induction of catabolic pathways to support the increased bioenergetics and biosynthetic needs of proliferative expansion, effector cytokine production and cytotoxicity(28). Several studies have shown that antigen-specific T cells upregulate the miR17∼92 cluster following activation to drive robust effector T cell responses(17–20, 22, 23, 34) (Fig S1A). However, the metabolic mechanisms by which miR-17∼92 drives effector T cell responses remains undefined.

To directly assess the function of miR-17∼92 in T cell metabolism, we engaged mouse models in which activation of CD4 and CD8 T cells results in conditional miR-17∼92 over-expression (conditional transgenic, CTg) or miR-17∼92 knock-out (conditional knockout, CKO) by using the human *GZMB* promoter to control Cre recombinase expression, and compared them to wild-type control mice (WT). This system leads to specific deletion or overexpression of miR17∼92 in mature peripheral T cells following activation, thus bypassing any developmental defects associated with loss of miR-17∼92 in germline deletion or transgenic models (18). We purified naïve WT, CTg and CKO T cells (Fig. S1B), which were subsequently activated *in vitro* with anti-CD3 and anti-CD28. Activated cells were then analyzed for their mitochondrial function by measuring the T cell oxygen consumption rates (OCR) and glycolysis via extracellular acidification rates (ECAR) in the presence of glucose as nutrient source by Seahorse analysis (Fig 1). Consistent with largely similar priming in WT, CTg and CKO T cells shown previously(18), similar upregulation in oxygen consumption rate (OCR) and glycolysis was noted in T cells following 24 hours of stimulation (Fig. S1C), likely due to similarly low levels of miR-17∼92 in WT, CKO and CTg cells during early stages of activation. However, 4 days after activation, when WT effector T cells express measurable levels of miR-17∼92, CKO effector CD4 T cells lacking miR-17∼92 were unable to upregulate oxidative phosphorylation to the levels of their WT and CTg counterparts as evidenced by reduced basal OCR and evident decrease in spare-respiratory capacity (SRC) (Fig. 1A). Furthermore, CKO CD4 T cells also demonstrated decreased glycolysis with a significant reduction in both ECAR as well as glucose uptake as measured by NBDG fluorescence (Fig. 1B, 1C). Consistent with these observations, gene set enrichment analysis (GSEA) demonstrated significant increase in the expression of mitochondrial respiration and oxidative phosphorylation gene signatures in miR-17∼92 expressing wild-type (WT) antigen-specific CD4 T cells compared to miR-17∼92 CKO CD4 T cells (Fig. S1D). Together these data implicate miR17∼92 as a critical driver of CD4 T cell glycolysis and mitochondrial respiration.

**Figure 1.**
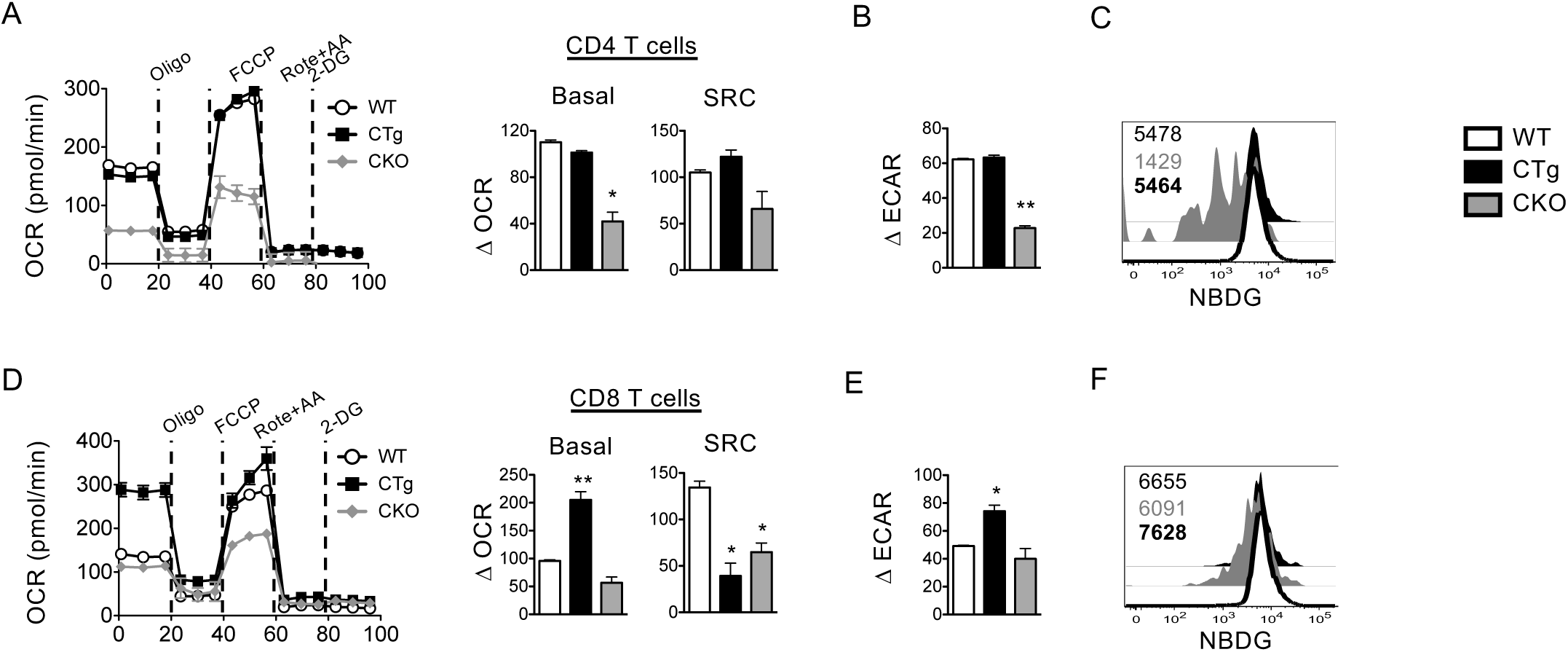
miR17∼92 drives glycolytic metabolism in activated T cells *in vitro*. WT, CTg, and CKO naïve mice were sorted for CD4 or CD8 T cells, which were then activated *in vitro* using αCD3+αCD28 for 4 days. CD4 and CD8 T cells were analyzed using XF Seahorse analyzer for glucose metabolism. **A**. Line graph depicts oxygen consumption rate of CD4 T cells after 4 days of *in vitro* activation. Bar graph shows basal OCR and SRC of WT, CTg and CKO CD4 T cells. **B**. Bar graph shows basal extracellular acidification rate (ECAR) at day 4 post activation. **C**. Histogram is gated on CD4 T cells showing NBDG uptake. D. Line graph depicts oxygen consumption rate of CD8 T cells after 4 days of *in vitro* activation. Bar graph shows basal OCR and SRC of WT, CTg and CKO CD8 T cells. **C**. Bar graph shows basal extracellular acidification rate at day 4 post activation. **D**. Histogram is gated on CD8 T cell showing NBDG uptake.

Ablation of miR-17∼92 in effector CD8 T cells resulted in a modest decrease in basal OCR and significant decrease in SRC, similar to CD4 T cells (Fig 1D). However, unlike CD4 T cells, which were unaffected by overexpression of miR-17∼92, CTg CD8 T cells exhibited a significant increase in basal oxidative phosphorylation upon overexpression of miR-17∼92 (Fig 1D). Intriguingly, the increase in basal OCR did not come with a corresponding increase in maximum OCR; such that CTg CD8 T cells showed significantly lower levels of SRC compared to the WT CD8 T cells (Fig 1D). These findings suggest that miR-17-92 does not affect the maximum mitochondrial capacity, but only alters the cell’s ability to utilize the mitochondria. Furthermore, over expression of miR-17∼92 in effector CD8 T cells increased the rates of aerobic glycolysis as demonstrated by the significantly higher basal ECAR in CTg compared to WT and CKO (Fig 1E). This increase in glycolysis was a function of glucose metabolism and not uptake, as there was no significant difference in the proportion of glucose uptake between the WT, CTg and CKO effector CD8 T cells (Fig. 1F). Following T cell priming effector CD4 T cells have been shown to be more dependent on oxidative phosphorylation pathways to meet their energetic demands, which is likely why the CKO phenotype is more pronounced when these cells fail to upregulate their mitochondrial machinery. In contrast, CD8 T cells are more dependent on glycolysis, which allows for miR-17∼92 over expression to significantly increase the less utilized oxidative pathway. Together these data implicate miR17∼92 as a critical driver of CD4 T cell glycolysis and mitochondrial respiration.

### miR-17∼92 drives effector T cell metabolic pathways through the AKT/mTOR axis

Based on the tumor suppressor PTEN being a direct target of miR-17∼92, we hypothesized that miR-17∼92 may support effector T cell expansion by promoting the downstream Akt/mTOR-dependent metabolic pathways. mTOR is the central nutrient sensor and metabolic regulator in T cells (35). Previous studies have demonstrated that PTEN is a direct target of miR-17∼92, and shows significant downregulation in CTg with a parallel increase in pS6 levels (Fig 2)(18, 20, 36). We next sought to determine whether the PTEN/Akt/mTOR pathway was acting downstream of miR-17∼92 in driving glycolysis in effector T cells. To evaluate this, WT, CTg and CKO T cells were stimulated *in vitro* in the presence or absence of the mTOR inhibitor rapamycin for 4 days, and then assessed for phosphorylation of S6 kinase target. We observed that CTg cells overexpressing miR-17∼92 were the most sensitive to rapamycin and showed almost a 4-fold decrease in pS6 MFI, while CKO cells were least sensitive with less than a 2-fold decrease in pS6 (Fig 2A). This decrease in pS6 levels directly correlated with a profound reduction in glycolysis in CTg effector cells as measured by a 3.8-fold change in basal ECAR levels (Fig 2B). Together these data demonstrated that miR-17∼92 affects T cell metabolism via mTOR upregulation, thus causing increased oxidative phosphorylation as well as glycolysis during effector expansion.

**Figure 2.**
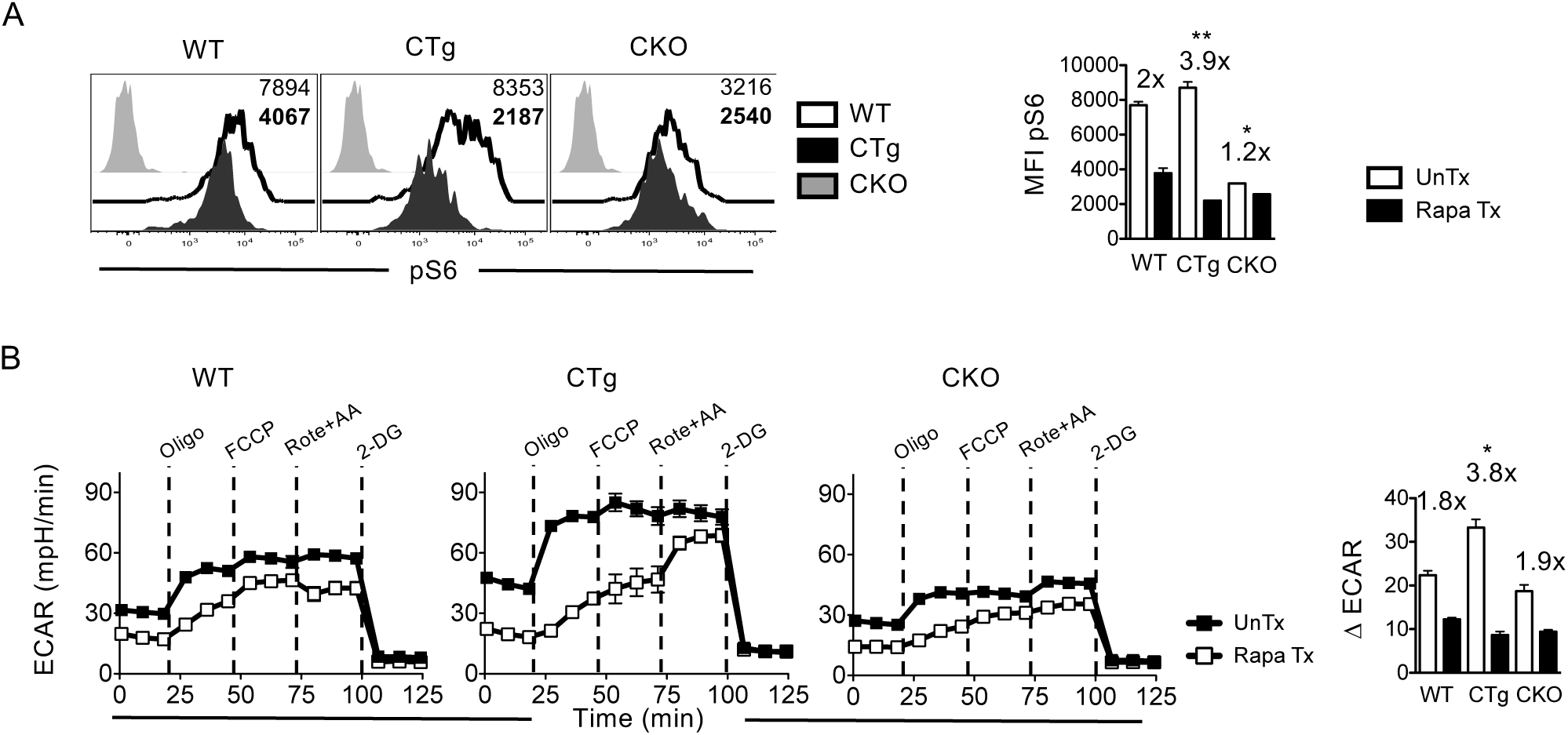
miR-17∼92 regulates T cell metabolism through the Akt/mTOR pathway. WT, CTg, and CKO CD8 T cells were purified and activated *in vitro* using αCD3+αCD28. Following 24 hours of priming CD8 T cells were treated with rapamycin and cultured for 3 days. CD8 T cells were stimulated using PMA+Ionomycin for 15 minutes. **A.** Histograms show pS6 levels in untreated (white) and Rapa Tx (Black) CD8 T cells. Bar graph shows MFI of pS6 in WT, CTg, and CKO CD8 T cells. **B**. Seahorse glucose assay was performed on CD8 T cells stimulated *in vitro* with and without Rapamycin. Line graphs show extracellular acidification rates (ECAR) at day 4 post-activation. Bar graph depicts basal ECAR at day 4 post-activation.

### miR-17∼92 is required during post-activation stages to drive expansion of antigen-specific CD4 and CD8 T cells

To better understand the T cell-intrinsic role of miR-17∼92 in driving the expansion of anti-viral T cell responses, we next analyzed the total numbers of antigen-experienced CD4 and CD8 T cells in WT, CKO and CTg mice following acute LCMV infection. Previous studies have engaged *CD4-Cre* regulated models of miR-17∼92 gene deletion or overexpression to query the role of miR-17∼92 in driving expansion and effector differentiation of T cells (6, 19, 22, 23). These models induce changes in miR-17∼92 expression during T cell development, which may result in defects in naïve T cells prior to activation. Hence, we leveraged the *GzmB-Cre* regulation model to specifically alter miR-17∼92 expression after T cell activation. In order to bypass any potential effects of differential expansion and effector differentiation of CKO or CTg T cells on viral clearance kinetics, we used the strategy of co-transferring WT LCMV specific SMARTA (GP61 specific TCR transgenic CD4 T cells) and P14 (GP33 specific TCR transgenic CD8 T cells) T cells to normalize viral clearance kinetics. Expansion of endogenous (WT, CKO or CTg) and donor (WT) CD4 and CD8 T cells was analyzed in LCMV-infected WT, CKO or CTg mice (Fig 3A).

**Figure 3.**
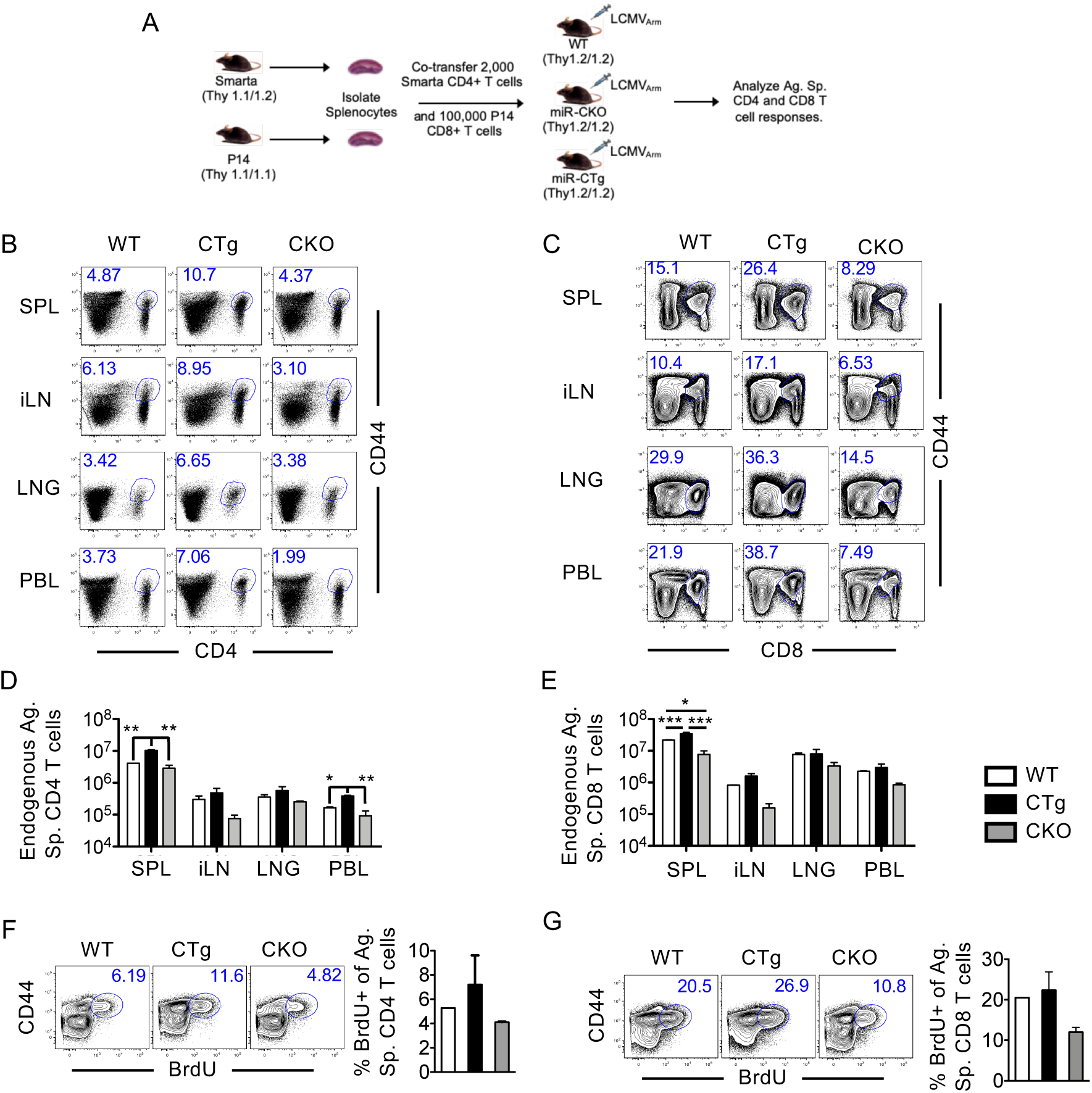
Induction of miR-17∼92 following activation promotes T cell expansion in a cell-intrinsic manner during viral infection. **(A)** Experimental setup. WT P14 and SMARTA cells were adoptively transferred into naïve WT, CTg, and CKO mice. Mice were subsequently infected with LCMV_Arm_ and tissues were analyzed at day 8 post-infection. (**B,C**) FACS plots show frequency of CD44+ CD4 T cells (**B**) and CD8 T cells (**C**) at day 8 post-infection. **D,E**. Bar graphs depict numbers of endogenous antigen-specific CD4 T cells (**D**) or CD8 T cells (**E**) in spleen, lymph node, lung and blood. **F,G**. FACS plots are gated on CD4+ (**F**) or CD8+(**G**) T cells. Numbers and bar graph depicts frequency of BrdU+ T cells. Data are representative of 2-5 experiments with n=3 mice per group. Bar graphs show mean and SEM. ANOVA with Tukey’s post-test was used with statistical significance in difference of means represented as * (P ≤ 0.05), ** (P ≤ 0.01), ***(P ≤ 0.001).

At the peak of anti-viral T cell responses (day 8 post-infection), over-expression of miR-17∼92 resulted in significant increase (2-fold) in total antigen-experienced CD44^Hi^ CD4 T in the spleens of CTg mice compared to WT hosts (Fig 3A, 3B). Similar, increases in endogenous antigen-specific CD4 T cells were also seen in other secondary lymphoid tissues (iLN) as well as peripheral sites such as PBMC and lung of CTg mice (Fig 3C). These differences were even more pronounced in the cytotoxic T cell responses with a significant increase in CD44^Hi^ CD8 T cells accumulating by day 8 post-infection following miR-17∼92 over-expression (Fig 3D). Conversely, genetic ablation of miR-17∼92 in CKO CD8 T cells led to stunted effector expansion compared to WT and CTg effector T cells (Fig 3A-D). Unlike the endogenous responses, donor CD4 and CD8 T cells accumulated in similar numbers by the peak of the LCMV response regardless of the miR expression in the recipient mice (Fig S2A, S2B), which suggests that the increased expansion observed in CTg effector T cells is a T cell intrinsic property of miR-17∼92 in driving effector expansion. Consistent with increased expansion, GSEA analysis of day 8 antigen-specific CD8 T cells from miR-17∼92 overexpressing CTg cells showed significant increase in mitosis gene signatures (Fig S2C). We next employed the strategy of *in vivo* administration of thymidine analogue, BrdU at 7.5 days post-infection to compare rate of S-phase progression. In the 12-hour window after injection, endogenous antigen-specific CD4 and CD8 T cells from CTg mice exhibited increased proliferation compared to WT mice as marked by greater BrdU incorporation (Fig 3F, 3G). In contrast, CKO mice exhibited moderately reduced BrdU incorporation compared to WT mice. These data demonstrate that miR-17∼92 expression following T cell activation is critical for driving the expansion of antigen-specific T cells by modulating CTL, and T_H_ cell proliferation.

### miR-17∼92 induction post-activation promotes the expansion of both T_H1_ and T_FH_ effector CD4 T cell lineages

We next assessed whether miR-17∼92 differentially regulated the expansion of virus-specific CD4 T cell responses (Fig 3) in a T_H_ lineage dependent manner. Previous studies using *Cd4*-Cre dependent deletion of miR-17∼92 in developing and naïve T cells prior to activation, implicated miR-17∼92 in T_FH_ and T_H1_ differentiation (6, 22, 23). As in Figure 3, in order to bypass any T cell developmental and T cell activation defects associated with loss or overexpression of miR-17∼92 prior to priming, we used the *GzmB*-Cre system to assess how changes in miR-17∼92 expression after CD4 T cell activation impact anti-viral T_H1_ and T_FH_ subsets following infection.

T_FH_ cells have been well characterized as CXCR5+, Ly6C-, T-bet^Lo^, Bcl-6^Hi^, and PD-1^Hi^ (37–41). Thus, we distinguished T_H1_ and T_FH_ responses in the I-A^b^GP61-specific CD4 T cells using Ly6C and CXCR5 (Fig 4A, 4B). This analysis revealed an evident increase in the proportions of T_H1_ cells in CTg mice compared to WT or CKO mice (Fig 4B), Detailed analysis of T_H1_ and T_FH_ cells with respect to lineage-specific transcription factors T-bet and Bcl-6 as well as cell surface marker PD-1 further confirmed the skewing of antigen-specific T_H_ responses towards T_H1_ subset upon miR-17∼92 over-expression (Fig 4C). As expected T_FH_ cells in WT, CTg, and CKO mice expressed higher levels of Bcl-6 and PD-1 and lower levels of T-bet compared to T_H1_ cells (Fig 4C). Thus, increased differentiation towards the T_H1_ is suggestive of either a decreased expansion of T_FH_ cells or an increased expansion of T_H1_ cells; to better understand this phenomenon we enumerated the T_H1_ and T_FH_ cells in secondary lymphoid and peripheral tissues. Following over-expression of miR-17∼92, we noted significantly increased numbers of both T_H1_ and T_FH_ cells in the spleen, and a moderate increase in T_H1_ responses in iLN and lungs (Fig 4D, 4E). A comparison of absolute numbers of T_H1_ and T_FH_ cells in CTg mice compared to WT mice showed about 2-3-fold increase in T_FH_ cell expansion upon over-expression of miR-17∼92 (Fig 4D, 4E). Notably, T_H1_ cells mounted 5-7-fold increased expansion following miR-17∼92 over-expression in both secondary lymphoid and peripheral tissues (Fig 4D, 4E).

**Figure 4.**
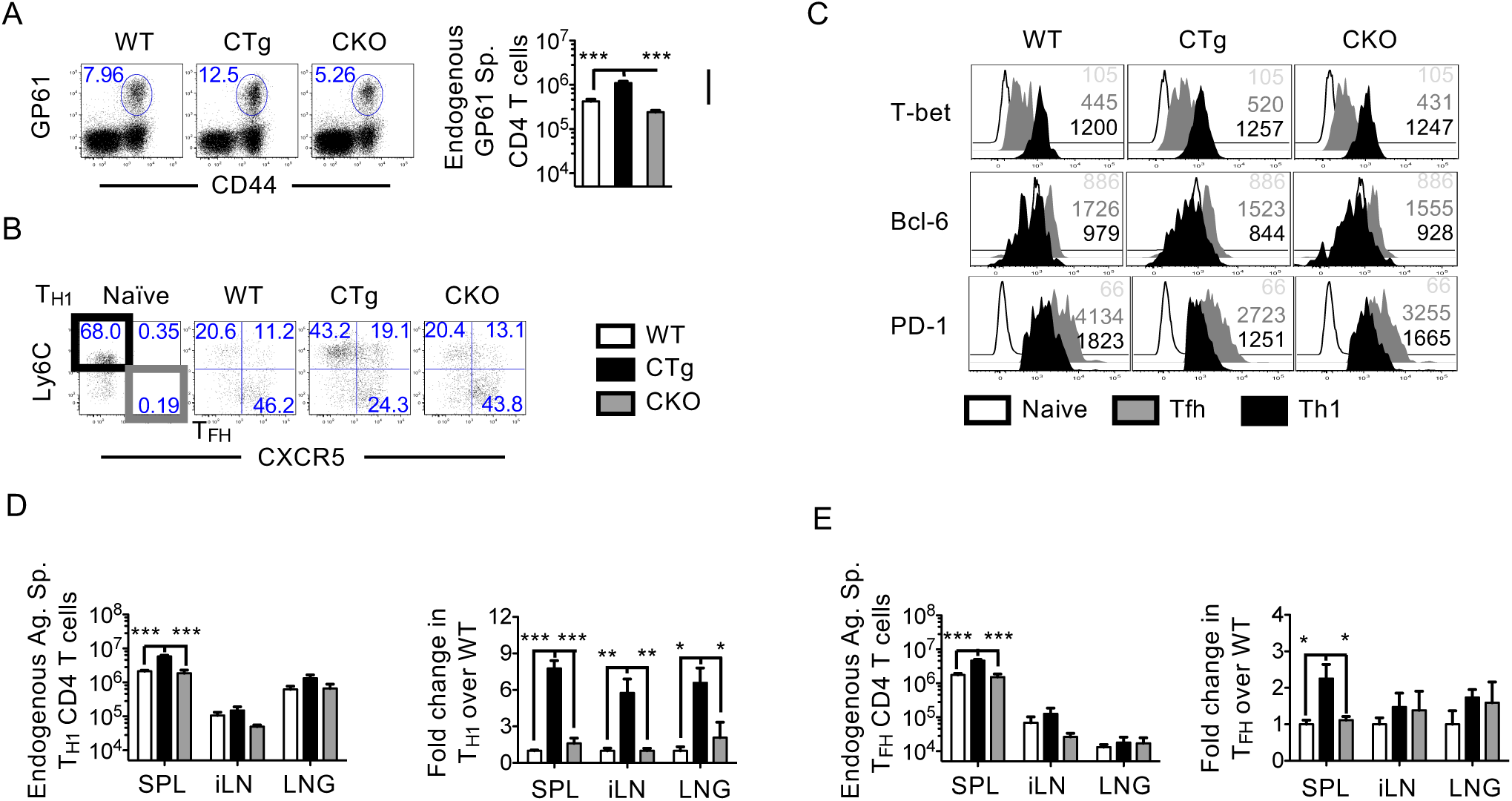
miR-17∼92 promotes development of T_H1_ and T_FH_ effector lineages in a T cell-intrinsic manner. WT P14 and SMARTA cells were adoptively transferred into naïve WT, CTg, and CKO mice. Mice were subsequently infected with LCMV_Arm_ and splenocytes were collected at day 8 post-infection. **A**. FACS plots are gated on CD4+ T cells. Number depicts frequency of GP61 specific endogenous CD8 T cells. Bar graph shows number of GP61 specific CD8 T cells in spleen. **B**. FACS plots are gated on endogenous GP61 specific CD4 T cells. **C**. Histograms are gated on GP61 specific CD4 T cells and then further sub-gated into T_H1_ and T_FH_ as shown in panel B. Number in histograms depict MFI of Naïve (light gray), T_FH_ (gray), and T_H1_ (black) CD4 T cells. **D-E**. Bar graphs depict number and fold-over WT of endogenous T_H1_ (**D**), and T_FH_ (**E**) GP61-specific CD4 T cells at day 8 post-infection in SPL, iLN, and LNG. Data are representative of at least 3 experiments with n=3 mice per group. Bar graphs show mean and SEM. ANOVA with Tukey’s post-test was used with statistical significance in difference of means represented as * (P ≤ 0.05), ** (P ≤ 0.01), ***(P ≤ 0.001).

Data in Fig 3 and S2 showed that increased CD4 and CD8 expansion following over-expression of miR-17∼92 is caused by intrinsic effects in effector T cells. Nonetheless, macrophages, NK cells, and CTLs also express granzyme B following activation and global immune dysregulation due to alterations in these cell-types upon miR-17∼92 over-expression or deletion could potentially cause CD4 T cell-extrinsic effects. Thus, to verify that the T_H1_ polarization we observed was CD4 T cell-intrinsic, we adoptively transferred congenically mismatched WT I-A^b^GP61-specific Smarta CD4 T cells prior to infection as in Fig 3. CD4 T cell differentiation events are critically regulated by the cytokine milieu, which could be different in WT, CTg and CKO mice. However, we observed that donor CD4 T cells had similar distributions of T_H1_ and T_FH_ responses in WT, CTg and CKO mice (Fig S3A-C), suggesting that global alteration in the inflammatory environment was not a major determinant of the T_H1_ polarization observed in endogenous CTg responses. Likewise, donor WT CTLs underwent similar effector differentiation, regardless of the environment, with similar expression of CD44, PD-1, T-bet and effector molecule GzmB (Fig. S3D). All together, these data demonstrate that miR-17∼92 drives CD4 T cell expansion of both T_H1_ and T_FH_ lineages in a T cell-intrinsic manner. While a previous study employing *miR-17∼92* gene deletion prior to T cell activation has implicated a specific role of miR-17∼92 in promoting T_FH_ differentiation (23), our data show a universal role of miR-17∼92 in driving the expansion of both T_H1_ and T_FH_ CD4 T cell lineages. Similar observations using CD4-Cre regulated miR-17∼92 expression (22) suggest that miR-17∼92 exerts minimal effects during T cell development or naïve T cell priming.

### miR-17∼92 drives terminal differentiation and loss of polyfunctionality in antigen-specific CD4 and CD8 T cells

Increased T cell expansion is often linked to increased terminal effector differentiation and a decrease in metabolic fitness (42–46). Our metabolism data showed that increased metabolic flux in CTg T cells paralleled the observed differences in effector T cell expansion in WT, CTg and CKO mice. Thus, we next sought to test whether the metabolic aberrancies were sustained through effector differentiation. To this end, we sorted CD44^Hi^ CD4+ and CD8+ effector cells at day 8 post-infection from spleens of infected animals and compared their metabolic states. Converse to their day 4 phenotype, CD4 T cells overexpressing miR-17∼92 had a significant drop in both basal metabolism, as well as the spare respiratory capacity (Fig 5A). CTg CD4 cells were also modestly defective in their glycolytic capacity (Fig. S4A, S4B). CTLs exhibited a similar decrease in basal oxygen consumption in endogenous CTg cells compared to WT control, as well as a significant drop in their SRC and basal ECAR (Fig 5B, S4C, S4D). Interestingly, CTg CD8 T cells displayed increased glucose uptake despite lower oxidative phosphorylation and glycolysis, which suggests that the energy yield is significantly reduced following effector expansion with miR-17∼92 over expression (Fig S4E). Furthermore, CD8 T cells in which miR-17∼92 was deleted following priming displayed a significant boost in both their spare respiratory capacity as well as their maximum ECAR, thus suggesting an increase in T cell fitness polyfunctionality and enrichment for MPEC phenotype (Fig 5A, 5B, S4A-D).

**Figure 5.**
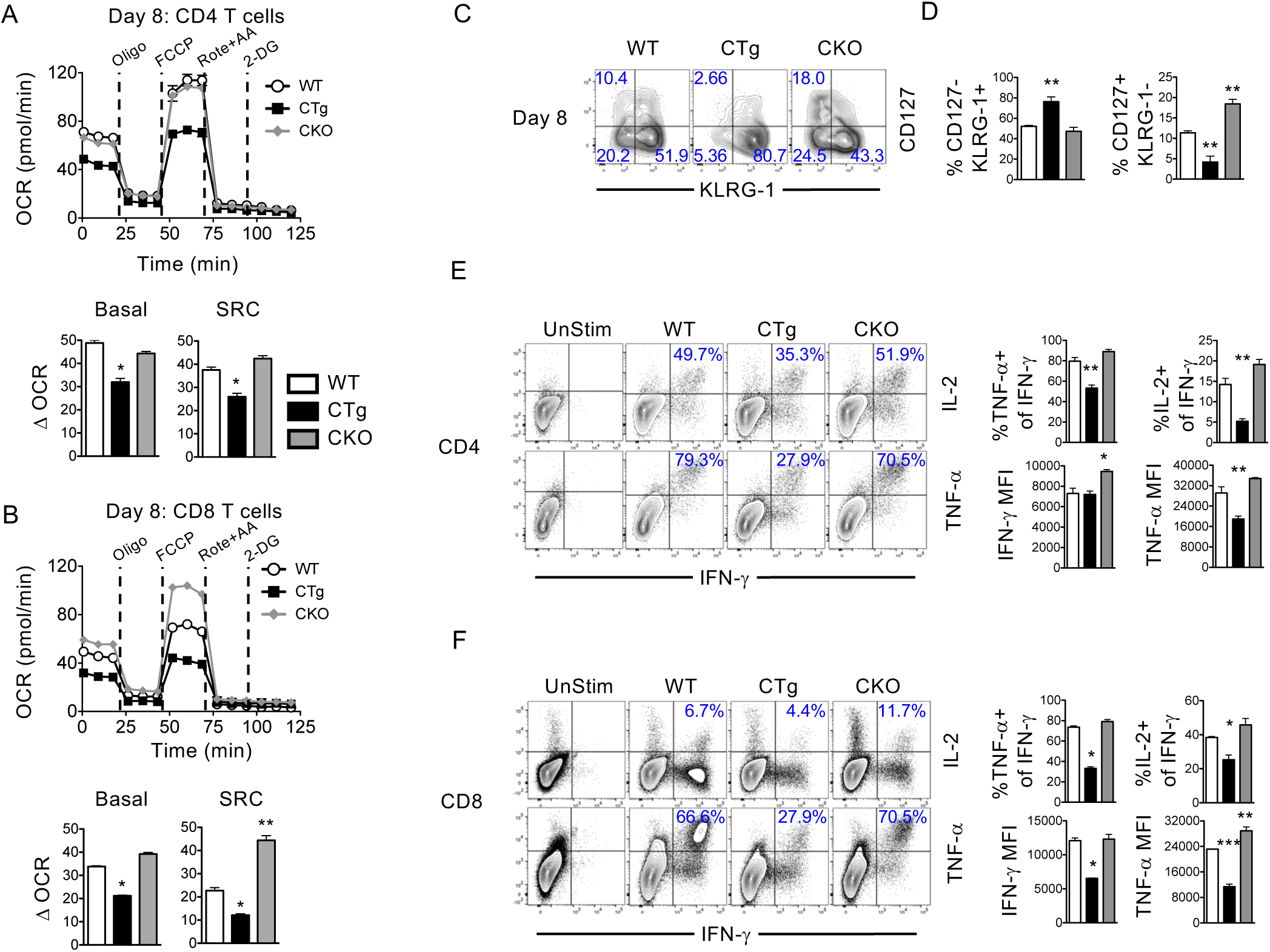
miR-17∼92 alters functional potency of T cells through metabolic dysregulation during viral infection. WT P14 and SMARTA cells were adoptively transferred into naïve WT, CTg, and CKO mice. Mice were subsequently infected with LCMV_Arm_ and splenocytes were collected at day 8 post-infection. **A-B**. WT, CTg, and CKO mice were infected with LCMV_Arm_ and splenocytes were collected at day 8 post-infection. Antigen-specific CD4+ (**A**) or CD8+ (**B**) T cells were purified and analyzed using a Mitostress test Seahorse assay. Line graphs show kinetics of oxygen consumption rates of T cells. Bar graphs show basal and spare respiratory capacity. **C**. FACS plots are gated on GP33+ CD8+ T cells. Numbers represent percent of antigen-specific CD8 T cells. **D**. Bar graphs depict frequency of CD127-KLRG-1+ (Terminal effectors) and CD127+KLRG-1-(Memory precursor effector cells). **E-F**. Splenocytes were stimulated with GP61 (**E**) or GP33 (**F**) peptides in the presence of BFA for 5 hrs and cytokine production was assessed. FACS plots are gated on CD4+ T cells (**E**) or CD8+ T cells (**F**). Frequency in plot depicts percent IL-2+ or TNF-α+ of IFN-γ+ CD4 T cells. Bar graphs show frequency of TNF-α+ or IL-2+ of IFN-γ+ and MFI of IFN-γ and TNF-α of IFN-γ+ T cells. Data are representative of 2-3 experiments with n=3 mice per group. Bar graphs show mean and SEM. ANOVA with Tukey’s post-test was used with statistical significance in difference of means represented as * (P ≤ 0.05), ** (P ≤ 0.01), ***(P ≤ 0.001).

Considering the stark decline in metabolic fitness observed in effector T cell differentiation following miR-17∼92 over-expression, we next sought to test if this affected the functionality of the effector T cells. To this end, antigen-specific T cells were analyzed for classical markers of memory T cell differentiation, and were stimulated at day 8 post-infection with LCMV specific cognate peptides gp61 (for antigen-specific CD4 T cells) and gp33 or gp276 (for antigen-specific CD8 T cells) to assess cytokine polyfunctionality. Over expression of miR-17∼92 resulted in a significant increase of terminal effector differentiation in GP33 specific CD8 T cells in CTg mice with a significant increase in KLRG-1+CD127-short-lived effector cell (SLEC) proportions, while CKO mice had an increased accumulation of CD127+KLRG-1-memory precursor effector cells (MPEC) (Fig 5C, 5D). These findings are consistent with the decrease in SRC observed in CTg and increase in CKO T cells, respectively.

Over expression of miR-17∼92 resulted in a significant decrease in the proportion of polyfunctional CD4 T cells with drop in TNF-α+IFN-γ+ and IL-2+IFN-γ+ CD4 T cell proportions (Fig 5E). Furthermore, when compared on a per cell basis, overexpression of miR-17∼92 had no effect on amounts of IL-2 produced, but caused a decrease in TNF-α production. However, deletion of this miR cluster caused an increase in IFN-γ production levels (Fig 5E, S4F). These results suggest an inverse relationship between miR-17∼92 expression and retention of T cell functional potency.

In line with the skewed MPEC and SLEC ratios in the cytotoxic T cell responses (Fig 5C, 5D), we observed a decrease in endogenous gp33 and gp276 specific polyfunctionality following miR-17∼92 over-expression as evidenced by a significant decrease in the proportions of TNF-α+ and IL-2+ of IFN-γ+ CD8 T cells (Fig 5F, S4H). Furthermore, CTLs from CTg mice produced significantly lower levels of IFN-γ and TNF-α compared to the WT controls with no significant effect on IL-2 production, while CKO were shown to be more functional with a higher production of TNF-α (Fig 5F, S4G, S4H). The fact that these differences were maintained for two distinct epitopes of LCMV further supports the point that these are T cell intrinsic effects and not caused by competition for antigen between donor and endogenous T cells.

## DISCUSSION

Research regarding the role of microRNA in the differentiation and function of CD4 and CD8 T cells have shown them to be key regulators of immunity. The wide role of CD4 T cells in regulation and modulation of systemic immunity and CD8 T cells in combating intracellular pathogens and tumors makes it especially important that we understand the roles that microRNAs play. This study demonstrates a unique role of miR-17∼92 in the modulation of T cell metabolism, thus resulting in increased expansion of CD8, T_H1_ and T_FH_ cellular immunity during anti-viral responses. Furthermore, we show that these differences are mTOR dependent, and that the unchecked metabolic activity related to miR17∼92 overexpression is ultimately associated with functional dysregulation. These findings support possible benefits to expressing miR-17∼92 in a controlled timely fashion so as to avoid T cell dysfunction in cancer immunotherapy approaches.

This study demonstrates for the first time that miR-17∼92 tunes the metabolic state of effector CD8 and CD4 T cells to a more glycolytic phenotype by upregulating glucose uptake in the newly primed effector cells with the concomitant increase in glycolysis and oxidative phosphorylation in a pS6-mTOR dependent manner. This was associated with increased expansion of T cells. CD4 and CD8 T cells vary in their extent of expansion following priming, with CTLs undergoing significantly greater expansion as seen by their number accumulation and BrdU uptake in WT cells. These differences suggest that the timing during which miR-17∼92 is critical for each T cell subset might be distinct. Despite these differences we observed that the miR-17∼92 cluster was critical in modulating early metabolic drive in both CD4 and CD8 T cells, which allowed for increased expansion of CTg T cells compared to WT and CKO cells. However, the continued utility of this metabolic program via miR-17∼92 overexpression drives terminal differentiation with loss of cytokine polyfunctionality and severe metabolic defect accumulation by the peak of the effector expansion, which reflected in both phenotypic and functional deficiencies.

Our findings are consistent with previous studies showing that miR-17∼92 exerts a role in driving antiviral CD8, T_H1_ and T_FH_ responses (22, 23). Unlike previous studies, which used CD4-Cre mediated changes in miR-17∼92 during thymic development, our studies show that miR-17∼92 driven expansion of CD8 T cells, T_H1_ and T_FH_ cells is a post-activation phenomenon, and not a function of aberrant T cell development. Furthermore, while T_FH_ expansion was also promoted by miR-17∼92, as previously reported (6, 22, 23), we observed a more profound impact on antiviral T_H1_ immunity, with almost twice as much expansion of T_H1_ cells compared to T_FH_ cells. It is possible that over-expression of miR-17∼92 during post-priming stages reduces the effect on the T_FH_ cells when compared to deletion during CD4 T cell development stages as published previously (22, 23). This notion is supported by the study by Kang *et al.*, where conditional miR-17∼92 deletion was achieved through OX40-regulated Cre expression (6); in this case also, the investigators observed a weaker phenotype with T_FH_ differentiation, possibly due to upregulation of OX-40 expression following activation, as in this study with GzmB-Cre. Collectively, these studies establish that miR-17∼92 exerts a definitive role in promoting metabolic conversion of CTL, T_H1_ and T_FH_ towards a more glycolytic state, strictly during the post-activation stages, and is not a manifestation of abnormal T cell development.

The wide-ranging impact of CD4 T helper and CTL functions in cellular and humoral responses make them important keystones of immunity; our study implicates miR-17∼92 as a potentially important target in optimizing these T cell responses during infections, immunizations and immunotherapies.

## ACKNOWLEDGMENTS

YY and FMB conducted experiments, analyzed data, interpreted the results, and prepared figures. YY also prepared the manuscript. SS and VK conceptualized the project, designed the experiments, supervised the work, analyzed data, interpreted the results, and prepared the manuscript. This work was supported by R01 award from NIAID to SS, R03 award from NIAID to VK, and seed funds from Seattle Children’s Research Institute to VK and SS. The authors would like to thank Ms. Shruti Bhise and Laura Penny for excellent technical assistance, and Dr. Joshy Jacob (Emory University, Atlanta) for kindly providing the *granzyme B-cre* mice.

## Supplemental Figures

**Supplemental Figure 1.**
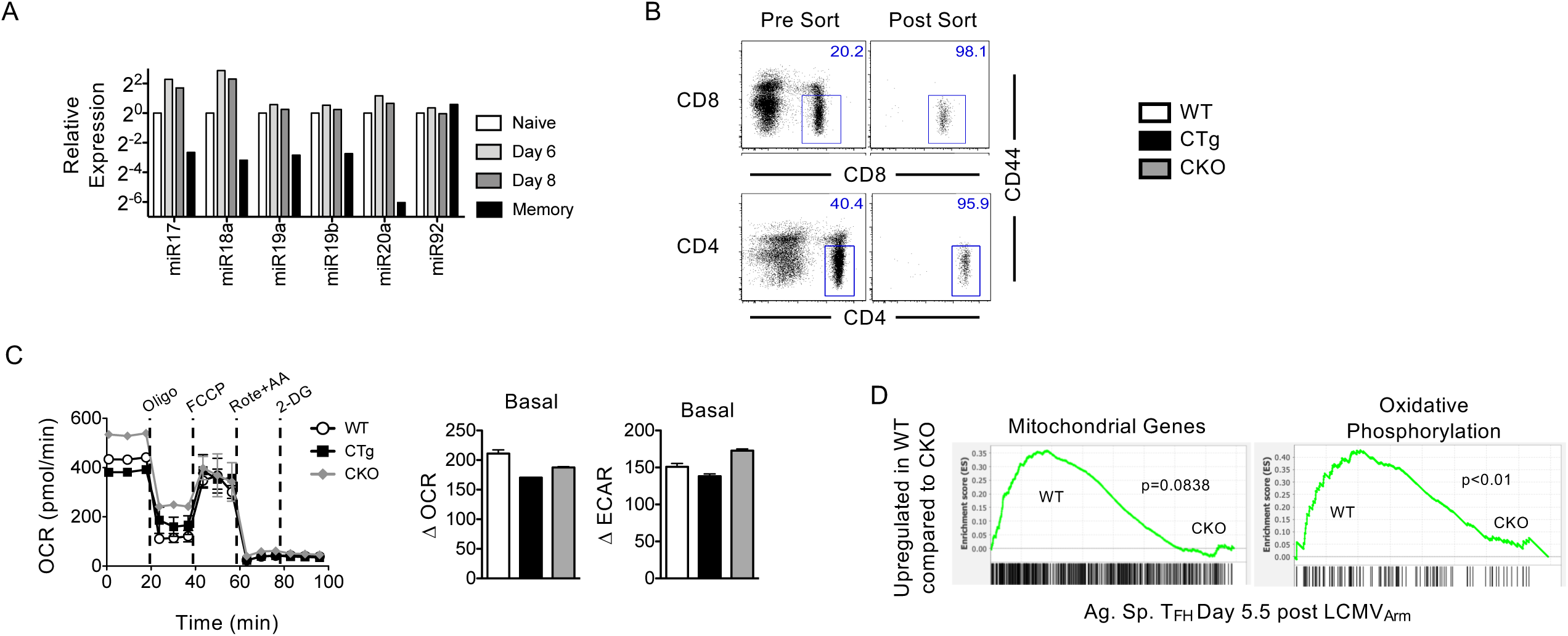
Dysregulation of CD8 T cell metabolic maintenance in miR over-expressing cells. (A) WT P14 T cells were adoptively transferred into naïve mice which were subsequently infected with LCMVArm. Splenocytes were collected at indicated times post-infection and donors were purified and miRNA isolated. Bar graph shows relative to naive expression of miR17∼92 in donors cells. (B) Naïve WT, CTg, and CKO splenocytes were isolated and CD4 and CD8 T cells were sort purified for CD4+ CD44lo or CD8+ CD44lo T cells. Representative pre- and post-sort data from WT group are shown. (C) Purified cells were activated in vitro using aCD3+aCD28. Line graph shows OCR of CD8 T cells at day 1 post-stimulation. C. Bar graphs depict basal OCR and ECAR of cells at day 1 post-stimulation. Data are representative of 2 experiments with n=3-5 mice per group. Bar graphs show mean and SEM. ANOVA with Tukey’s post-test was used with statistical significance in difference of means represented as ** (P ≤ 0.01). (D) Microarray data (Ansel et al.) at day 5.5 post-infection with WT and miR17∼92KO sorted TFH cells was used for GSEA analysis. GSEA plots are shown for Mootha Mitochondrial and Module 152 signatures looking at genes critical in mitochondrial maintenance and oxidative phosphorylation.

**Supplemental Figure 2.**
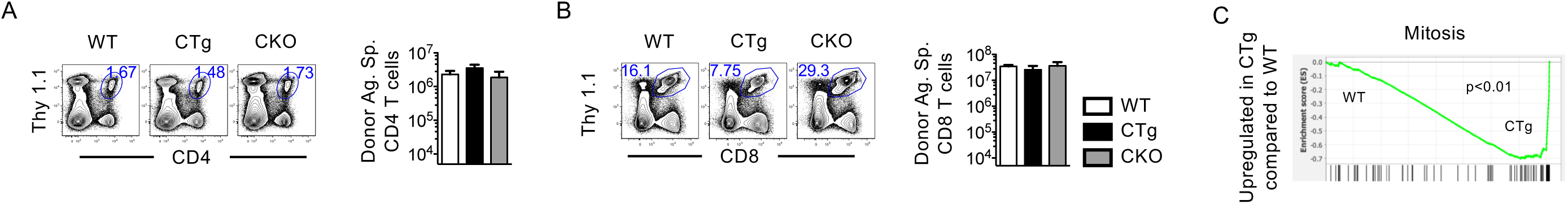
miR-17∼92 regulates T cell expansion via T cell intrinsic mechanisms. **B-C**. FACS plots show frequency of donor CD4 (**B**) or CD8 (**C**) T cells at day 8 post-infection. Bar graph depicts number of donor T cells in spleen. **D**. Microarray data (Wu et al.) at day 5.5 post-infection with WT and miR17∼92Tg sorted effector CD8 T cells was used for GSEA analysis. GSEA plots are shown for Mitosis gene signature. Data are representative of 2-5 experiments with n=3 mice per group. Bar graphs show mean and SEM. ANOVA with Tukey’s post-test was used with statistical significance in difference of means represented as * (P ≤ 0.05), ** (P ≤ 0.01).

**Supplemental Figure 3.**
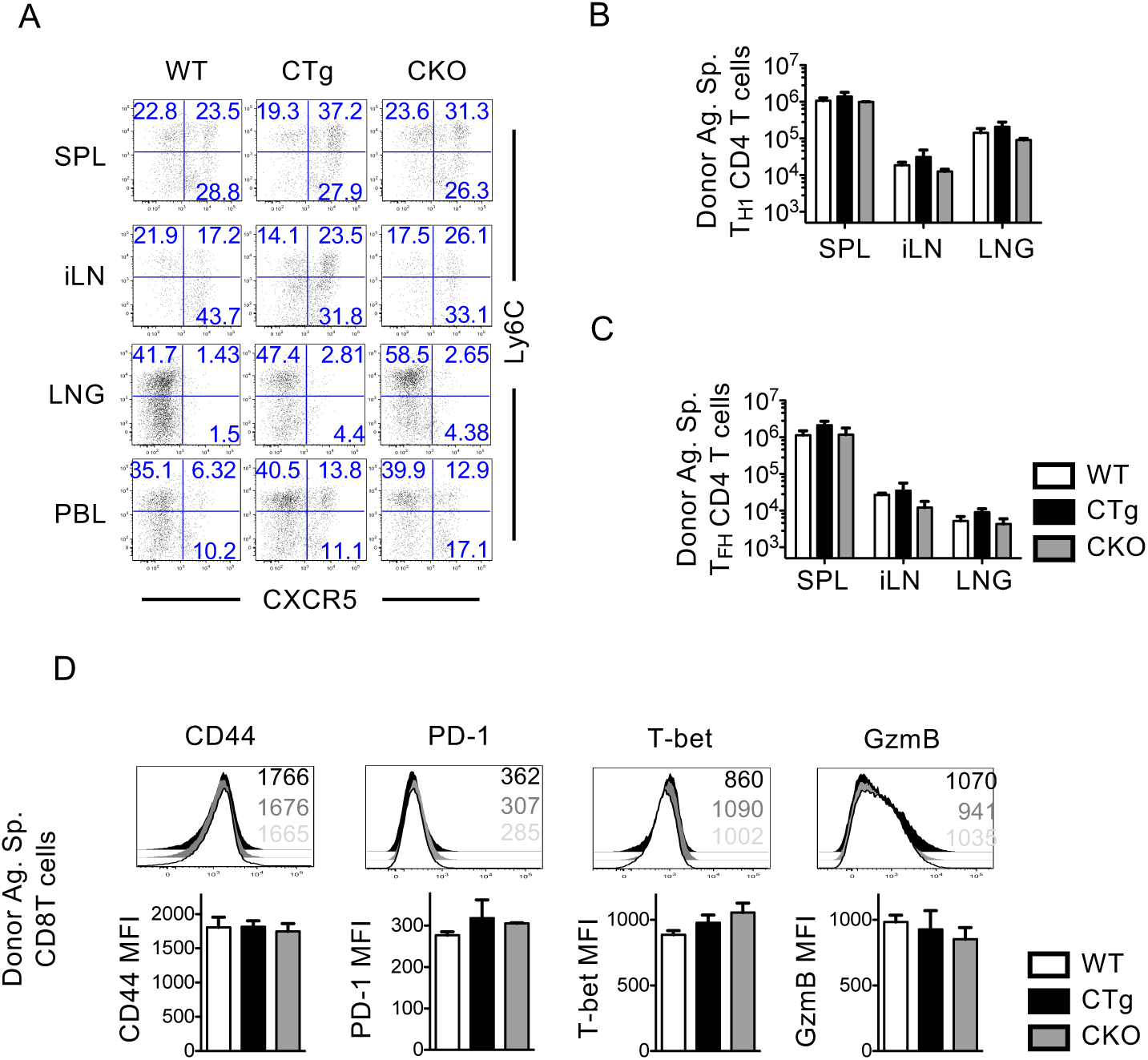
miR-17∼92 drives T cell intrinsic differentiation during acute infection. WT P14 and SMARTA cells were adoptively transferred into naïve WT, CTg, and CKO mice. Mice were subsequently infected with LCMVArm and splenocytes were collected at day 8 post-infection. **A**. FACS plots are gated on donor CD4+ T cells in SPL, iLN, LNG, and PBL. **B-C**. Bar graphs depict number of donor TH1 (**B**), and TFH (**C**) GP61-specific CD4 T cells at day 8 post-infection in SPL, iLN, and LNG. **D**. Histograms are gated on donor CD8 T cells. Numbers in histogram represent MFI of WT(light gray), CKO(gray), and CTg(black). Bar graphs show MFI of given markers in spleen. Data are representative at least 3 experiments with n=3 mice per group. Bar graphs show mean and SEM. ANOVA with Tukey’s post-test was used with statistical significance in difference of means represented as * (P ≤ 0.05), ** (P ≤ 0.01), ***(P ≤ 0.001).

**Supplemental Figure 4.**
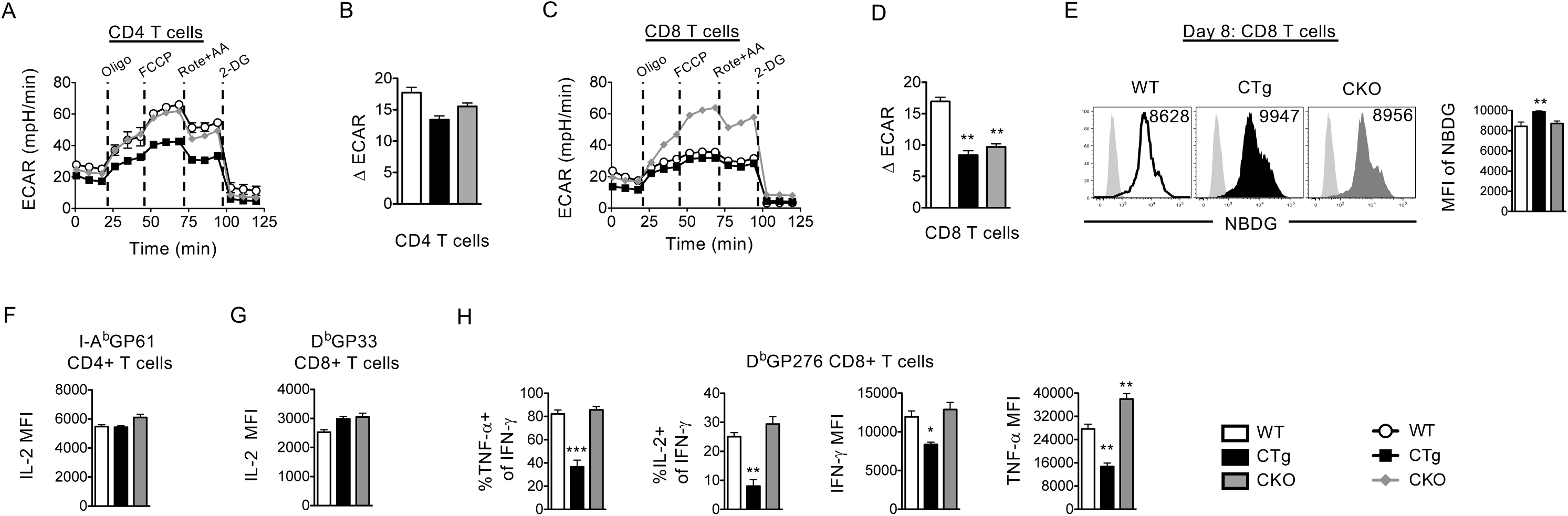
Metabolic and functional dysregulation upon constitutive overexpression of miR-17∼92. **A-D**. WT, CTg, and CKO mice were infected with LCMVArm and splenocytes were collected at day 8 post-infection. CD4+ (**A-B**) or CD8+ (**C-D**) T cells were purified and analyzed using a Mitostress test Seahorse assay. **A&C**. Line graphs show kinetics of oxygen consumption rates of T cells. **B&D**. Bar graphs show basal ECAR. **E**. Histograms are gated on CD8 T cells in day 8 splenocytes. Light-gray histogram show CD8 T cells untreated with NBDG. Number in histogram and bar graph show MFI of NBDG. **F-H**. Splenocytes were stimulated with GP61 (**F**), GP33 (**G**), or GP276 (**H**) in the presence of BFA for 5 hrs and cytokine production was accessed. Bar graphs show frequency of TNF-a+ or IL-2+ of IFN-g+ and MFI of IFN-g, TNF-a and IL-2 of IFN-g+ T cells. Data are representative of 2-3 experiments with n=3 mice per group. Bar graphs show mean and SEM. ANOVA with Tukey’s post-test was used with statistical significance in difference of means represented as * (P ≤ 0.05), ** (P ≤ 0.01), ***(P ≤ 0.001).

## REFERENCES

1. Jonas S, Izaurralde E. 2015. Towards a molecular understanding of microRNA-mediated gene silencing. Nat Rev Genet 16: 421–33

2. Gagnon JD, Ansel KM. 2019. MicroRNA regulation of CD8(+) T cell responses. Noncoding RNA Investig 3

3. Baumann FM, Yuzefpolskiy Y, Sarkar S, Kalia V. 2016. Dicer Regulates the Balance of Short-Lived Effector and Long-Lived Memory CD8 T Cell Lineages. PLoS One 11: e0162674

4. Liang Y, Pan HF, Ye DQ. 2015. microRNAs function in CD8+T cell biology. J Leukoc Biol 97: 487–97

5. Trifari S, Pipkin ME, Bandukwala HS, Aijo T, Bassein J, Chen R, Martinez GJ, Rao A. 2013. MicroRNA-directed program of cytotoxic CD8+ T-cell differentiation. Proc Natl Acad Sci U S A 110: 18608–13

6. Kang SG, Liu WH, Lu P, Jin HY, Lim HW, Shepherd J, Fremgen D, Verdin E, Oldstone MB, Qi H, Teijaro JR, Xiao C. 2013. MicroRNAs of the miR-17 approximately 92 family are critical regulators of T(FH) differentiation. Nat Immunol 14: 849–57

7. Jeker LT, Bluestone JA. 2013. MicroRNA regulation of T-cell differentiation and function. Immunol Rev 253: 65–81

8. Zhang N, Bevan MJ. 2010. Dicer controls CD8+ T-cell activation, migration, and survival. Proc Natl Acad Sci U S A 107: 21629–34

9. Baltimore D, Boldin MP, O’Connell RM, Rao DS, Taganov KD. 2008. MicroRNAs: new regulators of immune cell development and function. Nat Immunol 9: 839–45

10. Olive V, Jiang I, He L. 2010. mir-17-92, a cluster of miRNAs in the midst of the cancer network. Int J Biochem Cell Biol 42: 1348–54

11. Inomata M, Tagawa H, Guo YM, Kameoka Y, Takahashi N, Sawada K. 2009. MicroRNA-17-92 down-regulates expression of distinct targets in different B-cell lymphoma subtypes. Blood 113: 396–402

12. Mendell JT. 2008. miRiad roles for the miR-17-92 cluster in development and disease. Cell 133: 217–22

13. Xu W, Li JY. 2007. MicroRNA gene expression in malignant lymphoproliferative disorders. Chin Med J (Engl) 120: 996–9

14. Bui TV, Mendell JT. 2010. Myc: Maestro of MicroRNAs. Genes Cancer 1: 568–75

15. O’Donnell KA, Wentzel EA, Zeller KI, Dang CV, Mendell JT. 2005. c-Myc-regulated microRNAs modulate E2F1 expression. Nature 435: 839–43

16. Izreig S, Samborska B, Johnson RM, Sergushichev A, Ma EH, Lussier C, Loginicheva E, Donayo AO, Poffenberger MC, Sagan SM, Vincent EE, Artyomov MN, Duchaine TF, Jones RG. 2016. The miR-17 approximately 92 microRNA Cluster Is a Global Regulator of Tumor Metabolism. Cell Rep 16: 1915–28

17. Xiao C, Srinivasan L, Calado DP, Patterson HC, Zhang B, Wang J, Henderson JM, Kutok JL, Rajewsky K. 2008. Lymphoproliferative disease and autoimmunity in mice with increased miR-17-92 expression in lymphocytes. Nat Immunol 9: 405–14

18. Khan AA, Penny LA, Yuzefpolskiy Y, Sarkar S, Kalia V. 2013. MicroRNA-17∼92 regulates effector and memory CD8 T-cell fates by modulating proliferation in response to infections. Blood 121: 4473–83

19. Wu T, Wieland A, Araki K, Davis CW, Ye L, Hale JS, Ahmed R. 2012. Temporal expression of microRNA cluster miR-17-92 regulates effector and memory CD8+ T-cell differentiation. Proc Natl Acad Sci U S A 109: 9965–70

20. Jiang S, Li C, Olive V, Lykken E, Feng F, Sevilla J, Wan Y, He L, Li QJ. 2011. Molecular dissection of the miR-17-92 cluster’s critical dual roles in promoting Th1 responses and preventing inducible Treg differentiation. Blood 118: 5487–97

21. Wu Y, Heinrichs J, Bastian D, Fu J, Nguyen H, Schutt S, Liu Y, Jin J, Liu C, Li QJ, Xia C, Yu XZ. 2015. MicroRNA-17-92 controls T-cell responses in graft-versus-host disease and leukemia relapse in mice. Blood 126: 1314–23

22. Wu T, Wieland A, Lee J, Hale JS, Han JH, Xu X, Ahmed R. 2015. Cutting Edge: miR-17-92 Is Required for Both CD4 Th1 and T Follicular Helper Cell Responses during Viral Infection. J Immunol 195: 2515–9

23. Baumjohann D, Kageyama R, Clingan JM, Morar MM, Patel S, de Kouchkovsky D, Bannard O, Bluestone JA, Matloubian M, Ansel KM, Jeker LT. 2013. The microRNA cluster miR-17 approximately 92 promotes TFH cell differentiation and represses subset-inappropriate gene expression. Nat Immunol 14: 840–8

24. Baumjohann D, Ansel KM. 2013. MicroRNA-mediated regulation of T helper cell differentiation and plasticity. Nat Rev Immunol 13: 666–78

25. Vander Heiden MG, Cantley LC, Thompson CB. 2009. Understanding the Warburg effect: the metabolic requirements of cell proliferation. Science 324: 1029–33

26. Geltink RIK, Kyle RL, Pearce EL. 2018. Unraveling the Complex Interplay Between T Cell Metabolism and Function. Annu Rev Immunol 36: 461–88

27. Buck MD, Sowell RT, Kaech SM, Pearce EL. 2017. Metabolic Instruction of Immunity. Cell 169: 570–86

28. Sinclair LV, Cantrell DA. 2025. Protein Synthesis and Metabolism in T Cells. Annu Rev Immunol 43: 343–66

29. Jacob J, Baltimore D. 1999. Modelling T-cell memory by genetic marking of memory T cells in vivo. Nature 399: 593–7

30. Toumi R, Yuzefpolskiy Y, Vegaraju A, Xiao H, Smith KA, Sarkar S, Kalia V. 2022. Autocrine and paracrine IL-2 signals collaborate to regulate distinct phases of CD8 T cell memory. Cell Rep 39: 110632

31. Kalia V, Yuzefpolskiy Y, Vegaraju A, Xiao H, Baumann F, Jatav S, Church C, Prlic M, Jha A, Nghiem P, Riddell S, Sarkar S. 2021. Metabolic regulation by PD-1 signaling promotes long-lived quiescent CD8 T cell memory in mice. Sci Transl Med 13: eaba6006

32. Sarkar S, Yuzefpolskiy Y, Xiao H, Baumann FM, Yim S, Lee DJ, Schenten D, Kalia V. 2018. Programming of CD8 T Cell Quantity and Polyfunctionality by Direct IL-1 Signals. J Immunol 201: 3641–50

33. Kalia V, Penny LA, Yuzefpolskiy Y, Baumann FM, Sarkar S. 2015. Quiescence of Memory CD8(+) T Cells Is Mediated by Regulatory T Cells through Inhibitory Receptor CTLA-4. Immunity 42: 1116–29

34. Ohno M, Ohkuri T, Kosaka A, Tanahashi K, June CH, Natsume A, Okada H. 2013. Expression of miR-17-92 enhances anti-tumor activity of T-cells transduced with the anti-EGFRvIII chimeric antigen receptor in mice bearing human GBM xenografts. J Immunother Cancer 1: 21

35. Salmond RJ. 2018. mTOR Regulation of Glycolytic Metabolism in T Cells. Front Cell Dev Biol 6: 122

36. Simpson LJ, Patel S, Bhakta NR, Choy DF, Brightbill HD, Ren X, Wang Y, Pua HH, Baumjohann D, Montoya MM, Panduro M, Remedios KA, Huang X, Fahy JV, Arron JR, Woodruff PG, Ansel KM. 2014. A microRNA upregulated in asthma airway T cells promotes TH2 cytokine production. Nat Immunol 15: 1162–70

37. Oestreich KJ, Mohn SE, Weinmann AS. 2012. Molecular mechanisms that control the expression and activity of Bcl-6 in TH1 cells to regulate flexibility with a TFH-like gene profile. Nat Immunol 13: 405–11

38. Yu D, Rao S, Tsai LM, Lee SK, He Y, Sutcliffe EL, Srivastava M, Linterman M, Zheng L, Simpson N, Ellyard JI, Parish IA, Ma CS, Li QJ, Parish CR, Mackay CR, Vinuesa CG. 2009. The transcriptional repressor Bcl-6 directs T follicular helper cell lineage commitment. Immunity 31: 457–68

39. Crotty S. 2019. T Follicular Helper Cell Biology: A Decade of Discovery and Diseases. Immunity 50: 1132–48

40. Crotty S. 2021. Revealing T follicular helper cells with BCL6. Nat Rev Immunol 21: 616–7

41. Hale JS, Youngblood B, Latner DR, Mohammed AU, Ye L, Akondy RS, Wu T, Iyer SS, Ahmed R. 2013. Distinct memory CD4+ T cells with commitment to T follicular helper- and T helper 1-cell lineages are generated after acute viral infection. Immunity 38: 805–17

42. Buchholz VR, Flossdorf M, Hensel I, Kretschmer L, Weissbrich B, Graf P, Verschoor A, Schiemann M, Hofer T, Busch DH. 2013. Disparate individual fates compose robust CD8+ T cell immunity. Science 340: 630–5

43. Gerlach C, Rohr JC, Perie L, van Rooij N, van Heijst JW, Velds A, Urbanus J, Naik SH, Jacobs H, Beltman JB, de Boer RJ, Schumacher TN. 2013. Heterogeneous differentiation patterns of individual CD8+ T cells. Science 340: 635–9

44. Sarkar S, Teichgraber V, Kalia V, Polley A, Masopust D, Harrington LE, Ahmed R, Wherry EJ. 2007. Strength of stimulus and clonal competition impact the rate of memory CD8 T cell differentiation. J Immunol 179: 6704–14

45. Sarkar S, Kalia V, Haining WN, Konieczny BT, Subramaniam S, Ahmed R. 2008. Functional and genomic profiling of effector CD8 T cell subsets with distinct memory fates. J Exp Med 205: 625–40

46. Kalia V, Sarkar S, Subramaniam S, Haining WN, Smith KA, Ahmed R. 2010. Prolonged interleukin-2Ralpha expression on virus-specific CD8+ T cells favors terminal-effector differentiation in vivo. Immunity 32: 91–103

